# Recombinant venom proteins in insect seminal fluid reduces female lifespan

**DOI:** 10.1101/2024.01.14.575309

**Authors:** Samuel J. Beach, Maciej Maselko

## Abstract

The emergence of insecticide resistance has increased the need for alternative pest management tools^1,2^ Numerous genetic biocontrol approaches, which involve the release of genetically modified organisms to control pest populations, are in various stages of development to provide highly targeted pest control^3-7^. However, all current mating-based genetic biocontrol technologies function by releasing engineered males which skew sex-ratios or reduce offspring viability in subsequent generations. This allows mated females continue to cause harm (e.g. transmit disease). Here, we demonstrate the first example of *intra*generational genetic biocontrol, wherein mating with engineered males reduces female lifespan. The toxic male technique (TMT) involves the heterologous expression of insecticidal proteins within the male reproductive tract that are transferred to females via mating. We demonstrate TMT in *Drosophila melanogaster* males, which reduce the median lifespan of mated females by 37 - 59% compared to controls mated to wild type males. Agent-based models of *Aedes aegypti* predict that compared to existing genetic biocontrol technologies, even modest levels of mated female mortality could allow TMT to suppress a female population substantially faster, which is likely to result in reduced disease burdens. TMT presents a novel approach for combatting outbreaks of disease vectors and agricultural pests.

**Graphical Abstract:** 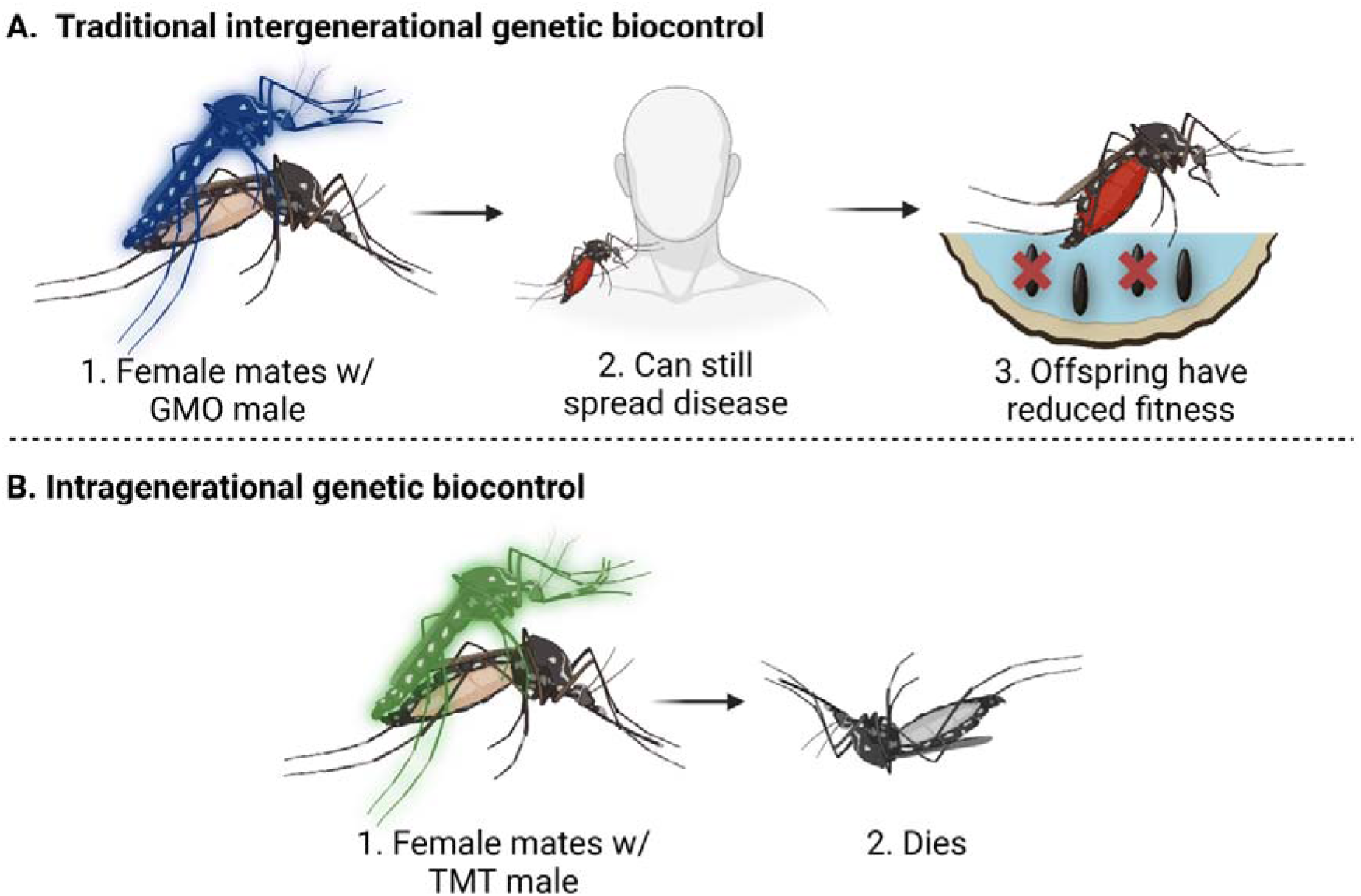

## Introduction

Insect pests pose a major challenge to human and environmental health. Malaria, spread by several species of *Anopheles* mosquitoes causes 608,000 deaths per year^8^ and rates of arboviral diseases spread primarily by the *Aedes aegypti* mosquito, including dengue, Zika, chikungunya, and yellow fever, are reaching unprecedented levels due to increased global trade and warming climates^9^. Dengue virus alone causes 390 million human infections each year and is now considered the most common vector-borne viral infection worldwide^10^. Recent studies have estimated that the cumulative cost of biological invasions is increasing four-fold every decade^11^, with a conservative estimate of US$162.7 billion for damages caused in 2017^12^. Invertebrate pests, primarily insects, are responsible for ∼40% of these damages^12,13^, and estimates of annual losses for all major food crops ranges between 20 – 30% due to damages by pests and pathogens^14^. Arboviral diseases such as avian malaria also pose a significant ecological threat to many native species, and are believed to be a key driver of population decline in New Zealand and Hawaiian forest birds^15,16^.

Pesticides are the first line of defence against many invasive species, particularly mosquitoes^17^. However, over-reliance on insecticides has resulted in widespread emergence of resistance^1^. Overapplication of insecticides has also resulted in declining populations of non-target species, and are responsible for many other environmental and human-health concerns^2^. Modern integrated pest-management is trending towards reducing reliance on chemical insecticides in favour of other environmentally friendly management techniques to reduce the emergence of resistances^1,17^.

Genetic biocontrol, defined as the release of organisms which have been genetically altered to reduce the spread and harm caused by a target species^18^, is one such alternative that’s been gaining broader public acceptance^19^. Early and well established genetic biocontrol technologies (GBT) such as the sterile insect technique (SIT) involve the release of mass numbers of males which have been radiologically sterilised^3^, thereby reducing the reproductive potential of the females which often only mate once in their lifetime. More modern GBT such as release of insects carrying a dominant lethal gene (RIDL)^4^ and gene drives^5^ function by propagating alleles which reduce the fitness of multiple future generations as the transgenes are inherited. These second generation GBT have proven to be more effective at supressing and eradicating pest populations in a shorter period with far fewer males required for release^20-22^ Currently, all GBT require a minimum of a generation to take effect on the target population and often much longer until the harm from a pest outbreak is mitigated^20^, with mated females continuing to blood feed and spread disease or damage crops by oviposition. Effects of female mating behaviour, such as commonly underestimated rates of polyandry^23^, have also been suggested to negatively affect the efficacy of mating-based GBT^24^. Current GBT are highly effective at long-term population control, but a more rapid response to outbreaks of disease vectors is required to reduce the spread of arboviruses and avert the risk of an epidemic^25^.

As endogenous seminal fluid proteins (SFP) are known to affect the physiology of mated females insects, including reducing mating receptivity and median lifespan^26,27^, we considered how insects may be genetically engineered so that their SFP can reduce female lifespan to a greater degree. SFP < 50 kDa have been found to pass through the female’s reproductive tract and into the haemolymph^28^, where they interact with targets accessible by the circulatory system. 60% of categorised venom compounds are cysteine-knotted mini-proteins^29^, which range from 2.6 – 14.8 kDa and are highly stable and proteolytic-resistant. Many of these venom proteins have been found to have highly specific activity to insect ion channels, with no toxicity found in mammals^30^

Here, we describe the toxic male technique (TMT), the first example of an intragenerational GBT. *Drosophila melanogaster were* engineered to heterologously express seven insect-specific venom proteins within the accessory glands of the male reproductive tract. When mated to wild type females, two of the TMT strains reduced median female lifespan by 37 – 59% compared to controls mated to wild type males. Single pair mating assays found that TMT males were able to court wild type females as effectively as wild type males. To determine the theoretical capability of TMT to suppress a pest population in comparison to female-specific RIDL (fsRIDL), the current state-of-the-art GBT, we developed an agent-based model to simulate an *Ae. aegypti field* trial^20^ Our model predicts that TMT should be capable of supressing a target female population substantially faster than fsRIDL, even at modest rates of mated female mortality. Importantly, our models also found that TMT reduced key epidemiological factors, such as cumulative bloodmeals and median number of gonotrophic cycles per female. These results demonstrate the potential of TMT as a new generation of genetic biocontrol which is uniquely suited to rapidly respond to outbreaks of disease vectors and agricultural pests.

## Results

### Heterologous expression of insecticidal venom proteins

We first identified a list of candidate venom proteins based on several criteria. Critically, the target of the venom could not be present in the male reproductive tract, which was determined by tissue specific gene enrichment values of the target ion channel given by Fly Atlas 2^31^ The venom proteins must also only interact with invertebrate ion channels, with no measurable LD50 found for mammals. Candidates were more likely to be chosen if they had lower Dipteran LD50, and if they had lower molecular weight, as we expect that to increase the likelihood of the venom proteins entering the haemolymph of mated females. To keep the number of venom proteins to be assessed manageable, we limited the list to the most promising candidate for a given target or venom class given our criteria (Table 1). Venom protein sequences were cloned into a UAS expression vector along with an *Acp26Aa* signal peptide to generate transgenic *D. melanogaster* strains.

**Table 1:** Candidate venom proteins investigated for accessory gland-specific recombinant expression.

| Venom protein | Native Host | Target |
| --- | --- | --- |
| μ-AGTX-Aa1d | *Agelenopsis aperta* | Na<sub>v</sub> channel (site 4) |
| Γ-CNTX-Pn1a | *Phoneutria nigriventer* | NMDA glutamate receptors |
| δ-CNTX-Pn1a | *Phoneutria nigriventer* | Na<sub>v</sub> channel (site 3) |
| κ-HXTX-Hv1c | *Hadronyche versuta* | BKCa channels |
| ω-HXTX-Hv1a | *Hadronyche versuta* | Ca<sub>v</sub>1 channels |
| ω-HXTX-Hv2a | *Hadronyche versuta* | Ca<sub>v</sub>2 channels |
| δ-AITX-Avd2a | *Anemonia sulcata* | Na<sub>v</sub> channel (site 3) |

### Testing functional expression of recombinant venoms

To assess the functionality of the heterologous venom proteins, homozygous UAS:venom strains were crossed to hemizygous *Act5C*-GAL4 flies^32^ that will either transmit the *Act5C*-GAL4 construct that will drive ubiquitous venom protein expression, or the CyO marker that results in a curled-wing phenotype. Progeny from strains functionally expressing venom proteins would only present the CyO phenotype, as inheritance of the *Act5C*-GAL4 chromosome would result in lethal ubiquitous expression of venom proteins. Conversely, progeny expressing nonfunctional venom proteins would exhibit 50% CyO phenotype, as ubiquitous expression would not decrease offspring fitness. From these crosses (N = 3), progeny from 5/7 of the UAS:venoms strains displayed 100% CyO phenotype, including μ-AGTX-Aa1d, Γ-CNTX-Pn1a, κ-HXTX-Hvlc, ω-HXTX-Hv1a, and δ-AITX-Avd2a. Progeny from UAS:venom strains expressing ω-CNTX-Pn1a exhibited 46±2% CyO phenotype, and co-HXTX-Hv2a exhibited 44±10%, so these venoms were presumed to be non-functional and not investigated further.

### Accessory-gland specific expression of transgenes

Success of this approach hinges on our ability to get high levels of accessory gland-specific expression of the venom proteins, while reducing off-target expression as much as possible. To determine the most suitable expression pattern, we tested three different accessory gland-specific GAL4 drivers: *antr*-GAL4; *prd*-GAL4; and *Acp26Aa*-GAL4^33-35^. These drivers were crossed to a UAS:*mCherry* reporter^36^, and the degree of on and off-target expression for each cross was determined by fluorescence microscopy of dissected male progeny (Fig. 1). Only *antr*-GAL4 produced a strong fluorescent signal in the accessory glands without also driving expression in earlier developmental stages or off-target tissue. Crosses of UAS:venom and *antr*-GAL4 strains produced equal numbers of male and female offspring which appeared to be healthy and fertile.

**Figure 1:**
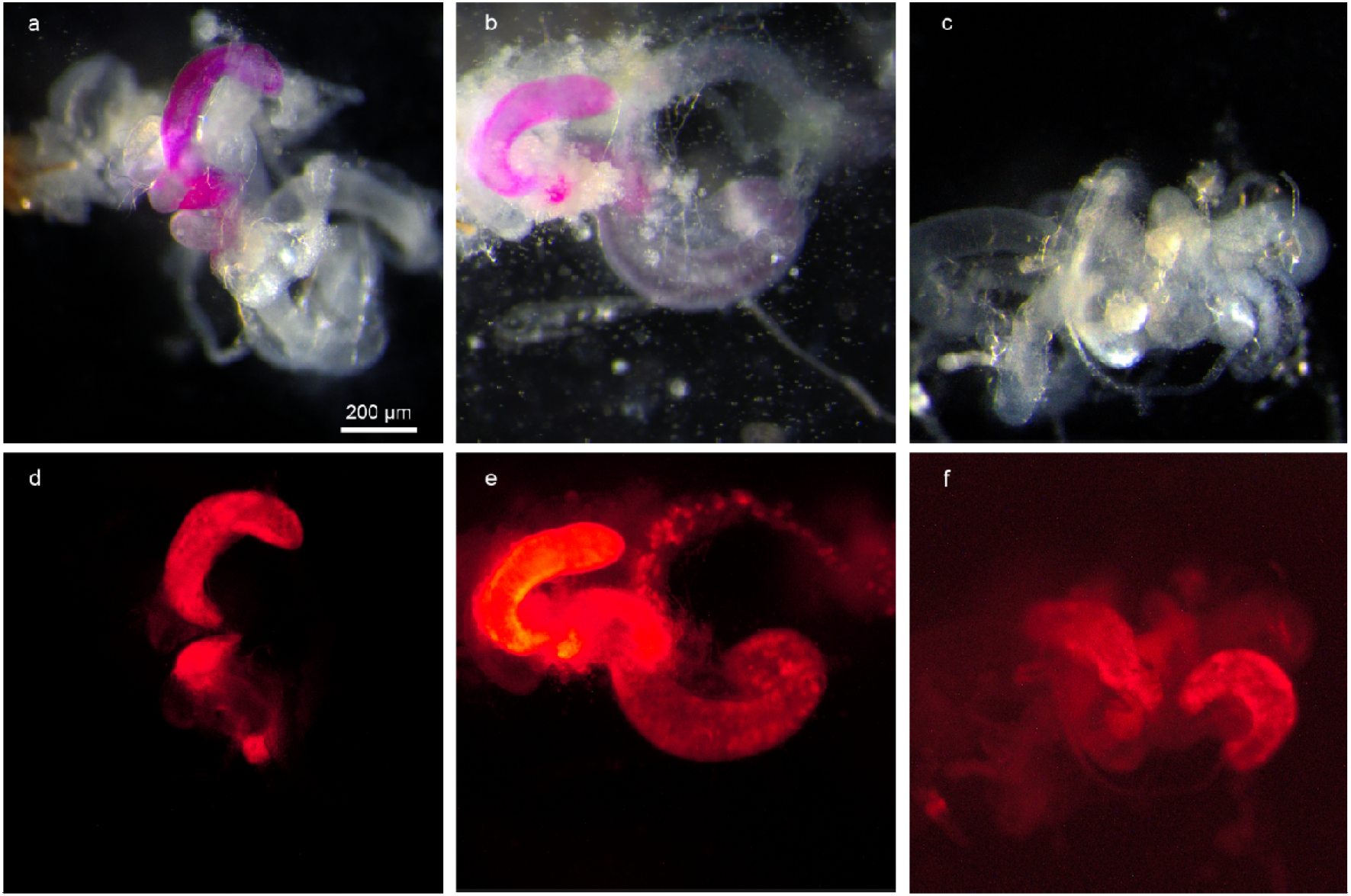
Dissected reproductive and digestive tracts of adult male offspring from UAS:*mCherry* reporter and accessory gland-specific GAL4 drivers (*antr*-GAL4 (a, d), *prd*-GAL4 (b, e), and *Acp26Aa-* GAL4 (c, f)) observed under bright-field (a-c) and red fluorescence (d-f) microscopy.

### TMT males reduce the lifespan of mated females

Mating assays were performed as previously described^37^ to determine whether TMT males can reduce the median lifespans of wild type females. TMT males were prepared by crossing UAS:venom strains (μ-AGTX-Aa1d, Γ-CNTX-Pn1a, κ-HXTX-Hv1c, ω-HXTX-Hv1a, or δ-AITX-Avd2a) to the accessory-gland specific *antr*-GAL4 driver strain, and transheterogous male offspring were isolated soon after eclosion. Virgin wild type females were housed either with an equal number of TMT males or wild type males and were transferred to fresh media every 3 days along with fresh males equal to the number of surviving females to maintain a 1:1 sex ratio. Females mated to wild type males had a median lifespan of 27 days, while females mated to T-CNTX-Pn1a TMT males had a 37% reduced median lifespan of 17 days (log-rank test, χ^2^ = 20.7, d.f. = 1, P = 5e-06; Fig. 2a), and females mated to δ-AITX-Avd2a TMT males had a 44% reduced median lifespan of 15 days (log-rank test, χ^2^ = 17.6, d.f. = 1, P = 3e-05 ; Fig. 2b). Females mated to μ-AGTX-Aa1d, κ-HXTX-Hv1c, or ω-HXTX-Hv1a TMT males did not have significantly reduced lifespans compared to wild type controls.

**Figure 2:**
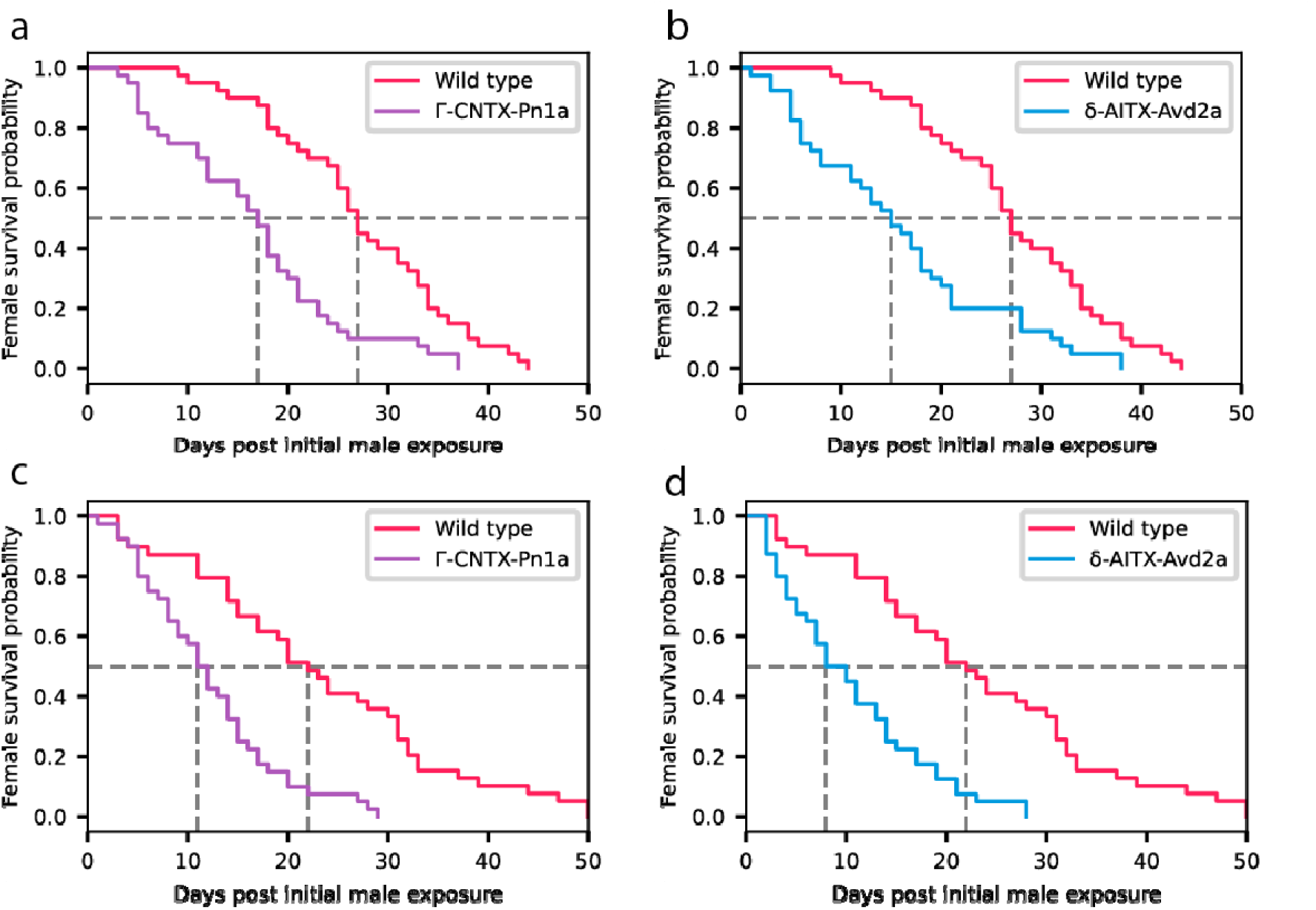
Survival probability of wild type female *D. melanogaster* mated to wild type males or TMT males expressing Γ-CNTX-Pn1a (a, c) or δ-AITX-Avd2a (b, d) venom proteins in their accessory glands. Females were exposed to males at an equal sex ratio (a, b), or with males at a three-fold excess (c, d).

Genetic biocontrol release programs utilise overwhelming ratios of modified males to wild individuals^20,38,39^ However, female *D. melanogaster* are known to have reduced fitness at higher rates of male exposure^40^. To determine if an increase in the male sex ratio of TMT males results in a greater than expected reduction in median lifespan, we performed another continuous mating assay with a 3:1 male-to-female sex ratio. In this assay, females mated to wild type males had a median lifespan of 22 days. Comparatively, females mated to Γ-CNTX-Pn1a TMT males had a 47% reduced median lifespan of 11.5 days (log-rank test, χ^2^ = 24.4, d.f. = 1, P = 8e-07; Fig. 2c), and females mated to δ-AITX-Avd2a TMT males had a 59% reduced median lifespan of 9 days (log-rank test, χ^2^ = 28.1, d.f. = 1, P = 1e-07; Fig. 2d). When compared to the 1:1 sex ratio mating assay, females mated to 3:1 wild type males did not have significantly reduced median lifespan (log-rank test, χ^2^ = 0.2, d.f. = 1, P = 0.6), while females mated to 3:1 Γ-CNTX-Pn1a and δ-AITX-Avd2a TMT males had a 32% (log-rank test, χ^2^ = 6.8, d.f. = 1, P = 0.009) and 40% (log-rank test, χ^2^ = 7.4, d.f. = 1, P = 0.007) respectively reduced lifespans relative to their equal ratio counterparts. This suggests that exposure to TMT males has dose-dependent effect on the lifespan of wild type females.

## TMT male fitness

### Single-pair courtship assay

The ability of a modified male released in a genetic biocontrol program to compete against wild males for access to mates is a critical component of the program’s performance^20^. To determine whether expression of venom proteins in the accessory glands impaired the males’ ability to court females, single-pair courtship assays were performed as previously described^41^. A single wild type virgin female was paired with a virgin male in a mating arena, which were observed for two hours to identify pairs which successful copulated. To control for off-target effects of accessory gland-specific expression of heterologous proteins, the competitiveness of negative control males expressing an unrelated low MW protein (NbVHH05^42^,13 kDa) in the same expression pattern (*antr*-GAL4/ UAS:*NBVHH05*) was also measured. The percentage of wild type males which successfully copulated was 30%, Γ-CNTX-Pn1a TMT males was 30%, δ-AITX-Avd2a TMT males was 22%, and negative control males was 24%. No significant difference was observed in the ability of the males to court females (χ^2^ = 1.3, d.f. = 3, P = 0.73).

### Competitive mating assay

Competitive mating assays, which assess the ability of genetically engineered males to compete with wild type males for mates, predominately utilize transgenic males prepared from wild type genetic background^6,7,43^. This is because males prepared using the GAL4-UAS system^44^, as well as ectopic expression of the *white (mini-w+)* marker, can confound the results of competitive mating assays due to differences in body size and colour^37^ or by increased rates of male-male courtship^45^. As synthetic accessory gland-specific promoters have yet to be characterized, the use of *antr*-GAL4 drivers were required to drive tissue-specific expression of the venom proteins. Characteristics unique to TMT, such as increased female mortality and differences in SFP composition, would likely further confound how male competitiveness is calculated by these assays (i.e. the proportion of transgenic offspring).

To determine if the inherent competitiveness of our GAL4-UAS transgenic males was negatively affected by variations in genetic background, we performed a control assay in which equal numbers of virgin wild type females, wild type males, and *antr-*GAL4/USA:*NbVHH05* males were housed together overnight and allowed to mate freely. Offspring were genotyped to determine paternity, and male competitiveness (C) was calculated as *C= P/(1 -P)*^*20*^, where P is the proportion of females which produced transgenic offspring, with *C = 1* indicating equal competitiveness to wild type males. The competitiveness of the control males was calculated to be just 0.14 (95% CI: 0.05-0.25), indicating that the GAL4-UAS background significantly affected female preference.

## TMT male longevity

Longevity assays were performed to determine what effect accessory gland-specific venom protein expression had on the longevity of TMT males relative to wild type and negative control males. Males were isolated soon after eclosion and housed in groups of 8, being transferred onto fresh media every 3 days. Γ-CNTX-Pn1a TMT males had no significant difference in longevity compared to wild type males (log-rank test, χ^2^ = 2.8, d.f. = 1, P = 0.09; Fig. 3) or negative control males (log-rank test, χ^2^ = 0.8, d.f. = 1, P = 0.4; Fig. 3). However, δ-AITX-Avd2a TMT males had 58% reduced median lifespan compared to wild type males (log-rank test, χ^2^ = 50-9, d.f. = 1, P = 1e-12; Fig. 3).

**Figure 3:**
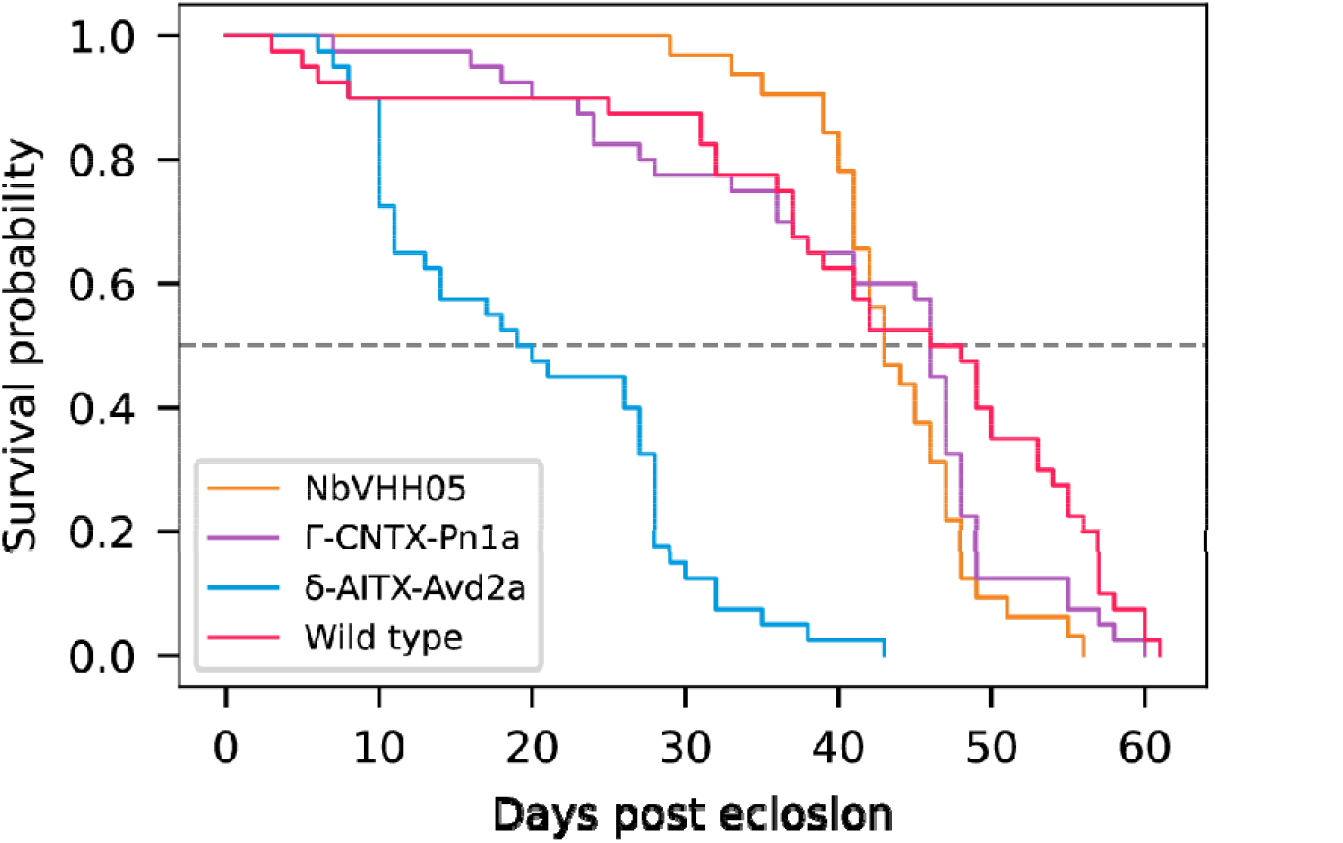
Longevity of TMT males compared to control strains. Γ-CNTX-Pn1a TMT males exhibit similar longevity to wild type and NbVHH05 negative control males, while δ-AITX-Avd2a TMT males show a marked decrease in survival.

## Modelling of *Ae. aegypti* population control

To evaluate how an TMT release program might compare to currently deployed GBT in its ability to suppress a population, we developed an agent-based model simulating *Ae. aegypti* field trials^20^ of SIT and fsRIDL mosquitoes. On average, the target female populations were reduced to 50% of their initial size (PR50) within 33.9 and 30.4 days respectively for SIT and fsRIDL release simulations, and were reduced by 95% (PR95) within 70 and 58.9 days respectively (Fig. 4a). For our model we assumed an idealized version of TMT and simulated the effect of female mating with TMT males to result in death (or inactivity ultimately leading to death) within an hour. The PR50 and PR95 of these TMT simulations were 12.7 and 39.2 days (Fig. 4a), 139% and 50% faster than fsRIDL respectively. The model also tracked key epidemiological factors such as cumulative blood feeding events, which were reduced on average by 61%, from 4295 in fsRIDL releases to just 1690 in TMT releases (Fig. 4b). The number of gonotrophic cycles correlates closely with two important factors, multiple feeding rates and the extrinsic incubation period (EIP) of arboviruses^46,47^, with the earliest that a female is likely to transmit a virus being her second gonotrophic cycle. The average number of subsequent (N > 1) gonotrophic cycles per female in fsRIDL and TMT releases was 3.84 and 1.82 respectively, a 53% reduction. We then investigated whether a median lifespan reduction of ∼50% as we’ve observed in our *D. melanogaster* assays would be sufficient to suppress or eliminate the target population. The PR50 of these ’prototype’ TMT releases was 22 days (Fig. 4a), 38% faster than fsRIDL. Cumulative blood feeding events and median gonotrophic cycles were also 27% lower in ’prototype’ TMT releases than fsRIDL, at 3142 and 2.78 respectively (Fig. 4b).

**Figure 4:**
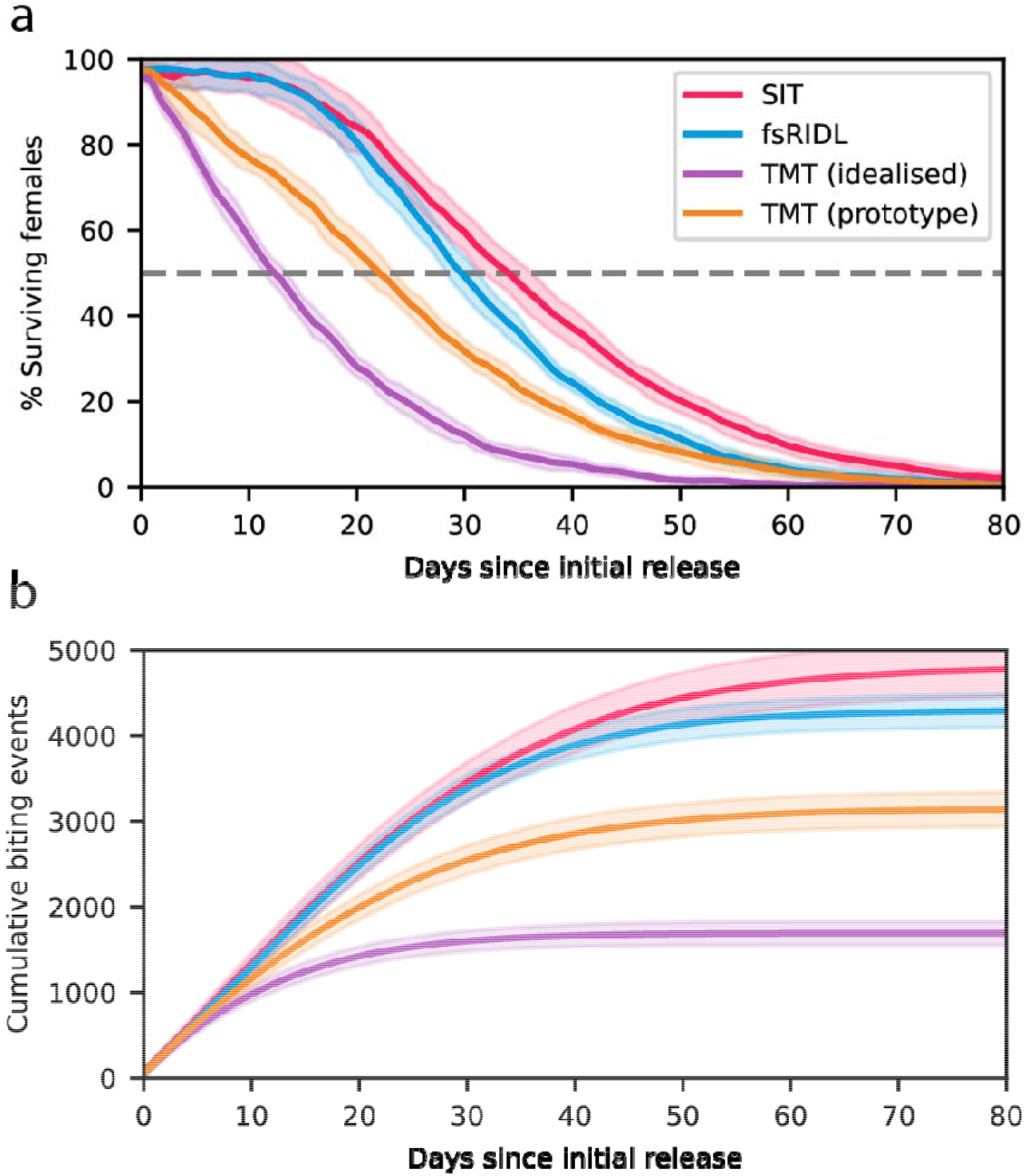
Agent-based models simulating field trials of *Ae. aegypti* population control programs, (a) The size of the female population during the release period relative to initial conditions. Data points are the mean of 10 simulations, the shaded area corresponds to the standard deviation, (b) The cumulative number of biting events which occurred throughout the releases. Data points are the mean of 10 simulations, the shaded area corresponds to the standard deviation.

## Discussion

Here, we describe the first example of a genetic biocontrol technology which can reduce the lifespan of mated females, called the toxic male technique, or TMT. The heterologous expression of insect-specific venom proteins, Γ-CNTX-Pn1a from the *P. nigriventer* spider and δ-AITX-Avd2a from the *A. sulcata* sea anemone, within the accessory glands of male *D. melanogaster* resulted in a 37–59% reduction in the median lifespan of mated wild type females. The development of intragenerational genetic biocontrol technologies such as TMT represents a paradigmatic shift in pest management, as their speed of population suppression may make them a viable alternative to chemical pesticides as a first line response to outbreaks, reducing off-target effects on local ecologies. TMT and comparable technologies could theoretically be implemented in any species where insemination occurs, provided that heterologous expression of the target protein can be constrained to the seminal fluid.

Our modelling suggests that even at modest rates of mated female mortality, TMT has the potential to suppress and eliminate female populations substantially faster than currently deployed GBT such as SIT and fsRIDL. These proven technologies are highly effective at local population suppression, but as they require at least a generation to take effect the mated females will persist in the target area and will continue to spread disease or damage crops. As female *Ae. aegypti* will most often mate prior to taking their first blood meal, females that mate with TMT males will be far less likely to transmit vector-borne diseases, with simulated TMT release programs reducing the incidence of blood feeding by 61% compared to fsRIDL releases. These results suggest that TMT may be uniquely well suited for rapidly responding to outbreaks of arboviral diseases, though further modelling will be required to understand the effects of epidemiology and other extrinsic factors (e.g. migration) on TMT’s efficacy.

The expression of Γ-CNTX-Pn1a in male accessory glands did not affect the lifespan of TMT males, but expression of δ-AITX-Avd2a reduced median lifespan by 58% compared to wild type males. We believe that this discrepancy is likely due to the difference in molecular weights of these venoms (5.2 kDa and 2.9 kDa respectively). The smaller size of δ-AITX-Avd2a may allow it to pass through the accessory gland epithelium into the haemolymph via paracellular routes, as has been described for some orally toxic insecticidal proteins^48^.The expression of either Γ-CNTX-Pn1a or δ-AITX-Avd2a did not reduce TMT male’s ability to court females in single-pair settings, though due to differences in genetic background we were unable to perform a meaningful assay to determine their ability to compete with wild type males for mates. Future iterations of TMT engineered in species such as *Ae. aegypti* will be prepared from strains of wild type genetic background, which will allow us to accurately assess their competitiveness in cage trial settings.

Over 1000 unique spider venom proteins have been characterised^30^, and many more from hosts such as scorpions and centipedes^49^. Many insecticidal venom proteins have toxicities that vary by orders of magnitude between different classes of insect^50^, which allows for targeted application of particular use cases. Co-expression of multiple toxins could help to mitigate the emergence of resistance alleles, as well as potentially resulting in synergistic toxicity from affecting multiple ion channel targets. Different combinations of venom proteins could be tailored for specific populations which have become resistant to insecticides or crops engineered to express Bt toxins. It needs to be determined whether heterologous venom protein expression may result in TMT males becoming toxic to natural predators. However, the oral toxicity of venom proteins is typically between 1 and 2 orders of magnitude lower than when they are directly injected^51^, and venom proteins can be selected which have significantly greater toxicity for the target species relative to natural predators.

Optimisations in the expression patterns of insecticidal proteins, such as changes to regulatory elements and bi-cistronic expression of multiple venoms, will likely result in further decreases in mated female lifespan. Dose-dependent responses to venoms tend to follow a sigmoidal curve, so depending on the current levels of expression it’s possible that even minor improvements could result in a significant increase in female mortality. Conditional expression or post-translational activation of venom proteins through mechanisms such as tetracycline-repression may also help to prevent fitness losses in TMT males. As UAS:venom/*antr*-GAL4 crosses produced fertile male and female offspring, it may be necessary to engineer TMT strains with genetic sorting of females by sex-specific intron splicing of venom expression, which would increase the efficiency of large-scale production of male-only cohorts and mitigate the risk of transgression of carrier female offspring.

## Methods

### Design and construction of UAS expression plasmids

The coding sequences for mature venom proteins (μ-AGTX-Aa1d, δ-CNTX-Pn1a, Γ-CNTX-Pn1a, κ-HXTX-Hv1c, ω-HXTX-Hv1a, ω-HXTX-Hv2a, δ-AITX-Avd2a), NbVHHOS, as well as the *Acp26Aa* signal peptide, were codon optimised for expression in *D. melanogaster* using Geneious Prime 2023.2.1. pJFRC81-10XUAS:IVS-Syn21-GFP-p10 (hereafter pJFRC81) was a gift from Gerald Rubin (Addgene plasmid #36432 ; http://n2t.net/addgene:36432 ; RRID:Addgene_36432). The region 67 bp upstream of the start codon of the GFP CDS of pJFRC81 was used as an extended upstream homology arm to re-introduce the Syn21 5’UTR during assembly. 30 bp of the p10 *3’* UTR was used as the downstream homology arm, with a Sapl recognition site introduced between the stop codon and terminator for future mutagenesis. Sequences comprised of the *Acp26Aa* signal peptide, a coding sequence, and both homology arms were ordered as gBlocks from Integrated DNA Technologies (IDT) and assembled (NEBuilder® HiFi DNA Assembly Master Mix (New England Biolabs (NEB))) into pJFRC81 after linearisation with Notl and Xbal. All plasmids were sequence verified by Plasmidsaurus (Eugene, OR) whole plasmid sequencing. Plasmid DNA sequences are listed in Supplementary Table S1.

### Fly strains, transgenesis, and husbandry

Canton S wild type flies were sourced from the Bloomington Drosophila Stock Centre (RRID: BDSC #64349). Act5C-GAL4/CyO flies were sourced from the Bloomington Drosophila Stock Centre (RRID: BDSC #4414). UAS:*mCherry* flies were sourced from the Bloomington Drosophila Stock Centre (RRID: BDSC #52268). *prd-GAL4, antr-GAL4*, and *Acp26Aa-GAL4* flies were a kind gift from Mariana F. Wolfner from Cornell University.

UAS expression plasmids were sent to BestGene Inc (Chino Hills, CA) for plasmid DNA midipreps, microinjections into a strain with a 2nd chromosome attP docking site for ϕC31 mediated genomic integration (RRID: BDSC #9752), and outcrossing of transgenic lines to balancer strains. Homozygous UAS stocks were prepared by selecting against the CyO marker on the balancer chromosome.

Flies were maintained on cornmeal diet based on the Bloomington Drosophila Stock Centre (BDSC) standard Nutri-Fly formulation (catalogue number 66-113; Genesee Scientific). Flies were reared in a controlled environment room at 25°C, 75% humidity, and a 12 h light-dark cycle with a 30 min transition period. Biosafety approval was granted by the Macquarie University Institutional Biosafety Committee (#5202).

### Recombinant venom functionality assay

To test the functionality of recombinant venom expression, hemizygous *Act5C*-GAL4/CyO virgin females were crossed to homozygous UAS:(μ-AGTX-Aa1d, δ-CNTX-Pn1a, Γ-CNTX-Pn1a, κ-HXTX-Hv1c, io-HXTX-Hv1a, œ-HXTX-Hv2a, δ-AITX-Avd2a) males. Three crosses were prepared for each of the venoms, with each cross comprising of 10 females and 10 males which mated freely for 7 days before the parents were discarded. Offspring were collected for up to 10 days after the parents were discarded, and the presence or absence of the CyO marker was scored for each offspring.

### Characterising accessory gland-specific GAl4 drivers

The strength of on and off-target heterologous gene regulation by various accessory gland GAL4 drivers was determined by crossing UAS: *mCherry* reporter strains to *prd-GALA, antr-GAL4*, and *Acp26Aa* -GAL4 driver strains. Offspring were observed throughout their development, and adult male offspring were dissected to isolate their reproductive tracts. The strength of off-target and accessory gland-specific expression of mCherry was identified by fluorescence microscopy (Olympus SZX16 stereomicroscope, 6.3x magnification, Excitation: 530-550 nm, Emission: 575 nm longpass).

## Lifespan assays

### Continual mating assays

Wild type females were isolated soon after eclosion and housed together in groups of eight for 2 - 3 days prior to male exposure to ensure that they would be responsive to male courtship. Transheterozygous antr-GAL4/UAS:(μ-AGTX-Aa1d, T-CNTX-Pn1a, κ-HXTX-Hv1c, œ-HXTX-Hvla, δ-AITX- Avd2a) males were prepared continuously throughout the experiment by crossing homozygous UAS males to homozygous ontr-GAL4 virgin females. Male offspring, including wild type males, were isolated soon after eclosion and housed in groups of eight for 3 - 5 days prior to female exposure. Five groups of eight females were allocated to each treatment group. Females were briefly anaesthetised with CO2 to be transferred to fresh vials every 3 days, and deaths were scored every 1 - 2 days. Males were initially added at this first transfer (2-3 days post-female eclosion) and were replaced during subsequent transfers to maintain the desired sex ratio, as well as to ensure that females had constant access to robust males with full reserves of SFP.

### Male lifespan assays

Transheterozygous *antr*-GAL4/UAS:(r-CNTX-Pn1a, δ-AITX-Avd2a, NbVHH05) males were prepared by crossing homozygous UAS males to homozygous ontr-GAL4 virgin females. 40 male offspring, including wild type males, were isolated soon after eclosion and housed in five groups of eight. Males were briefly anaesthetised with CO2 to be transferred to fresh vials every 3 days, and deaths were scored every 1–2 days.

## Mating assays

### Single-pair courtship assays

200 virgin wild type females, 50 virgin wild type males, and 50 virgin transheterozygous *antr-* GAL4/UAS:(Γ-CNTX-Pn1a, δ-AITX-Avd2a, NbVHH05) males were isolated soon after eclosion and housed in groups of 10–15. Males were aged for 3 – 5 days, and females for 2 – 3 days prior to the experiment. Mating arenas were prepared by adding 6.2 g of sucrose diet (5% sucrose, 2% agar) to 35 mm petri dishes, leaving enough head room for the flies to comfortably move around.

Experiments were performed at the same time of day, starting 30 minutes after dawn. Flies were briefly anaesthetised, and single pairs were transferred to mating arenas. The pairs were observed for 2 hours, successful copulations were recorded, and the fraction of successful copulations was calculated to be the percentage courtship of a given treatment group.

### Competitive mating assay

50 virgin wild type females, 50 virgin wild type males, and 50 virgin *antr*-GAL4/UAS:NbVHH05 males were isolated soon after eclosion and housed in groups of 5. Males were aged for 3 - 5 days, and females for 2 - 3 days prior to the experiment. Assays were prepared just before the dusk to reduce visual courtship stimuli and increase reliance on chemosensory cues^52^. Flies were briefly anaesthetised, and ten groups of five wild type females, five wild type males, and five transgenic males were transferred to fresh media. The flies were immediately placed into darkened environment chambers and were allowed to mate freely until the next morning, at which point the males were discarded and the females were isolated on fresh media. 10 offspring from each female was collected, genomic DNA was extracted from pooled offspring as previously described^53^, which was then used as template for 20 μL PCR reactions with OneTaq® DNA Polymerase (New England Biolabs (NEB)). The quality of the extracted genomic DNA was determined by using primers against the endogenous *white* locus using the following primers: white-F, and white-R (Supplementary Table S2). Paternity of the offspring was determined by the presence or absence of the UAS:NbVHH05 locus using the following primers: UAS:F, and UAS:R (Supplementary Table S2).

### Agent-based modelling

To model the expected performance of TMT relative to existing GBT such as fsRIDL, we developed an agent-based model of Ae. *aegypti* using GAMA 1.9 (https://gama-platform.org). The model is not spatially explicit, but instead assumes a homogenous, randomly mixing population, of a size for which the major limiting factor is L1-L2 instar larval density-dependent mortality. We incorporated empirically derived parameters such as density-independent daily survivorship, durations of immature stages, and length of gonotrophic cycles (all parameters and sources listed in Supplementary Table S3). Rates of density-dependent larval mortality were tuned such that the population equilibrium fits observations from field studies^54^, with ∼15% of the total agents in the population being adult mosquitos. 10 simulations were run for each treatment, each instantiated with 2500 eggs, with hourly time steps following the progression of each individual throughout their life stages (egg, L1/L2 instar larvae, L3/L4 instar larvae, pupae, adult) until the population reached a steady state of ∼ 500 adult mosquitos. Adults are assumed to be inactive at night and the middle of the day, and during active periods females will either find a mate, blood feed, lay eggs, or rest depending on their needs at a given cycle. Mating lethality (the likelihood of a mate dying within that cycle) for the idealized iteration of TMT releases was set to 100%. Mating lethality for ’prototype’ TMT males was calibrated to 55%, which was derived by simulating releases with a 3:1 release ratio, with releases every three days and 100% male competitiveness, then tuning the lethality such that the median female lifespan of the target population was reduced from 17 days to 9 days (47% reduction). Females that mate with fsRIDL males will produce heterozygous offspring with 100% late-stage female mortality, females that mate with SIT males will produce no offspring, and females that mate with but are not killed by ’prototype’ TMT males will produce wild type offspring. Parameters for transgenic male release were modelled after the suppression phase of the Brazil field trials of OX513A mosquitoes^20^, including values for transgenic male competitiveness (3.1%), release ratio (25:1 transgenic males to total wild population), and release period (every 3 days).

### Statistical analysis

Lifespans of mated females and transheterozygous males was analysed by log-rank statistical analysis using R 4.3.2 (R Core Team, 2023), the *survival* (v3.5-7; Terry M Therneau, 2023) and the *survminer* (v0.4.9; Alboukadel Kassambara, 2021) packages. To test for differences in single-pair male courtship, a one-way chi-square test for contingency tables were used to calculate P values. Confidence intervals for the mating competitiveness of *antr*-GAL4/UAS:NbVHH05 males were calculated using the Wilson method.

## Figures

Kaplan-Meier survival curves and cumulative biting rates of model *Ae. aegypti* populations were plotted using Matplotlib^55^ in Python 3.11.5. Graphical abstract was prepared using BioRender.

## Data availability

All data are available in the main text or the supplementary materials.

## Code availability

GAMA 1.9 model files are available via the Maselko Lab GitHub (https://github.com/MaselkoLab/).

## Acknowledgements

The authors are grateful to Prof. Mariana F. Wolfner from Cornell University for providing many of the GAL4 strains used in this project, and to Prof. Glenn King from the University of Queensland for sharing his expertise on venom proteins and their heterologous expression.

## Author Information

### Contributions

S.J.B. and M.M. conceived this study. S.J.B. and M.M. designed experiments and analysed data. S.J.B. performed the experiments and developed the model. All authors contributed to writing the manuscript.

### Corresponding author

Correspondence to Maciej Maselko

